# The Lettuce Expression Browser: from lab to LEB

**DOI:** 10.1101/2025.10.24.684171

**Authors:** Dirk-Jan M. van Workum, Esther S. van den Bergh, Siddhant S. Shetty, Jelmer van Lieshout, Tao Feng, Marrit C. Alderkamp, Gema Flores Andaluz, Alan Pauls, Flip F. M. Mulder, Dmitry Lapin, Mark G. M. Aarts, Guido van den Ackerveken, Remko Offringa, Ronald Pierik, Marcel Proveniers, M. Eric Schranz, Sandra Smit, Renze Heidstra, Anneke Horstman

## Abstract

Lettuce (*Lactuca sativa* L.) is an economically important leafy vegetable within the Asteraceae family, cultivated worldwide across diverse agricultural systems. Recent advances in genomic and transcriptomic resources have positioned lettuce as a promising model system for functional genomics in the Asteraceae. However, currently available gene expression datasets lack comprehensive tissue-specific resolution, primarily focus on a single cultivar and are not visualised in an interpretable manner, limiting their utility for broader genetic and physiological studies. To bridge this gap, we developed the Lettuce Expression Browser (LEB), a publicly available platform providing high-resolution gene expression maps across various organs, tissues and developmental stages in both cultivated and wild lettuce species. The LEB integrates transcriptomic data from finely dissected seedlings, shoot tissues at various developmental stages and seedlings subjected to abiotic stresses (salt and far-red), visualised using the ggPlantmap R package. This platform offers an intuitive interface for exploring gene expression patterns and serves as a valuable resource for those studying lettuce development, stress responses, and evolutionary genomics. The LEB is hosted on the LettuceKnow Web Portal (https://lettuce.bioinformatics.nl) and can be expanded to include additional datasets, enhancing its role as a key tool for lettuce research and crop improvement.

**Significance statement:** The Lettuce Expression Browser (LEB) provides the first high-resolution gene expression atlas for both cultivated and wild lettuce species. This open-access resource enables detailed exploration of gene activity across development stages and stress conditions, advancing functional genomics in the Asteraceae family.

## Introduction

Lettuce (*Lactuca sativa* L.), a major leafy vegetable crop in the Compositae (Asteraceae) family, is cultivated and consumed worldwide. With a production value of approximately 20 billion USD in 2020 (FAOSTAT, 2023), lettuce holds significant economic importance and is highly adaptable to modern farming methods such as vertical farming and hydroponics. Believed to have been domesticated from *Lactuca serriola* over 6,000 years ago, lettuce has since diversified into a range of types, including leaf types (cos, butterhead, crisp) and non-leaf types (stalk and oilseed). Distinct domestication traits, such as leaf morphology and loss of seed shattering, separate cultivated lettuce from its close wild relatives. These include *L. serriola*, *L. saligna* and *L. virosa*, which provide valuable breeding traits, such as disease resistance and stress tolerance (Zhang et al. 2017; Wei et al. 2021).

Lettuce has recently gained recognition as a promising model system for the Asteraceae family, largely due to the growing availability of genomic and transcriptomic resources. To fully establish lettuce as model species, high-resolution, easily accessible gene expression data across different cultivars are essential in addition to genomic sequences. Comprehensive gene expression profiles spanning developmental stages, organs and tissues, and environmental treatments are crucial for uncovering underlying molecular mechanisms. Browsable gene expression data enable this next level of insight, similar to those available for well-established model species such as *Arabidopsis thaliana*, and will greatly enhance our understanding of lettuce biology.

The publication of the first reference genome for lettuce of the crisphead cultivar ‘Salinas’, a large-scale RNA-sequencing study in 2017 and a major population genomics study in 2021 have significantly advanced lettuce research (Reyes-Chin-Wo et al. 2017; Wei et al. 2021; Zhang et al. 2017). With the development of genome editing technologies like CRISPR/Cas9, lettuce has become an increasingly useful species for functional genomics research (Bertier et al. 2018). Existing resources, such as the Lettuce Genome Database (http://www.lettucegdb.com/) (Guo et al. 2023) and the Lettuce Database (https://db.cngb.org/lettuce/) (Zhou et al. 2024), have aggregated these datasets and added germplasm and phenotype data, along with some expression data. However, these resources are limited by poor tissue resolution and focus on a single cultivar, making them far from comprehensive. This underscores the need for more detailed tissue- and organ-specific gene expression data, as exemplified by expression browsers for model species like *Arabidopsis thaliana* (Winter et al. 2007), as well as for major crops like tomato and maize (Stelpflug et al. 2016; Hoopes et al. 2019; Rose 2016; Fernandez-Pozo et al. 2017; Waese et al. 2017).

To address this, we have developed the Lettuce Expression Browser (LEB), which includes a detailed tissue-specific gene expression map and a developmental series, both of which cover a range of tissues in both cultivated and wild lettuce species. Additional datasets include *L. sativa* shoot (leaf) and whole root samples under various conditions (control, salt and supplemental far-red light exposure). The gene expression data are visualised through the recently developed ggPlantmap R package (Jo and Kajala 2024), which uses colour gradients to display expression levels across individual plant parts. This setup mirrors the widely used electronic Fluorescent Pictograph (eFP) Browser (Waese et al. 2017), providing a familiar and intuitive interface for plant biologists.

The Lettuce Expression Browser, hosted on the LettuceKnow Web Portal (https://lettuce.bioinformatics.nl), provides easy access to the data and can be expanded as new datasets become available. This initiative sets out to create an invaluable tool for studying context-dependent, spatially resolved gene expression across both cultivated and wild lettuce species. Future updates will include a broader array of tissues, developmental stages, and diverse lettuce accessions, establishing it as a critical resource for lettuce research and crop improvement.

## Results

To aid in the interpretation and analysis of transcriptomic data across diverse lettuce organs and tissues, treatments, developmental stages, and accessions, we developed the Lettuce Expression Browser (LEB), available at https://lettuce.bioinformatics.nl/leb/. This user-friendly tool overlays gene expression levels onto intuitive, graphical representations of experimentally collected samples. The LEB enables researchers to explore spatial and temporal gene expression patterns across multiple conditions and genotypes in both cultivated and wild lettuce. Below, we describe the datasets used to build the LEB and provide examples of its application for studying tissue-specific and stress-responsive gene expression.

### Description of samples

We sampled accessions of four *Lactuca* species, including several lettuce cultivars, across different growth substrates, growth conditions and organs or tissues. This generated four distinct gene expression maps (**Figure 1**). By incorporating finely dissected plants, developmental stages and important abiotic stresses, we provide a comprehensive overview of gene expression relevant to major aspects of lettuce biology.

**Figure 1:**
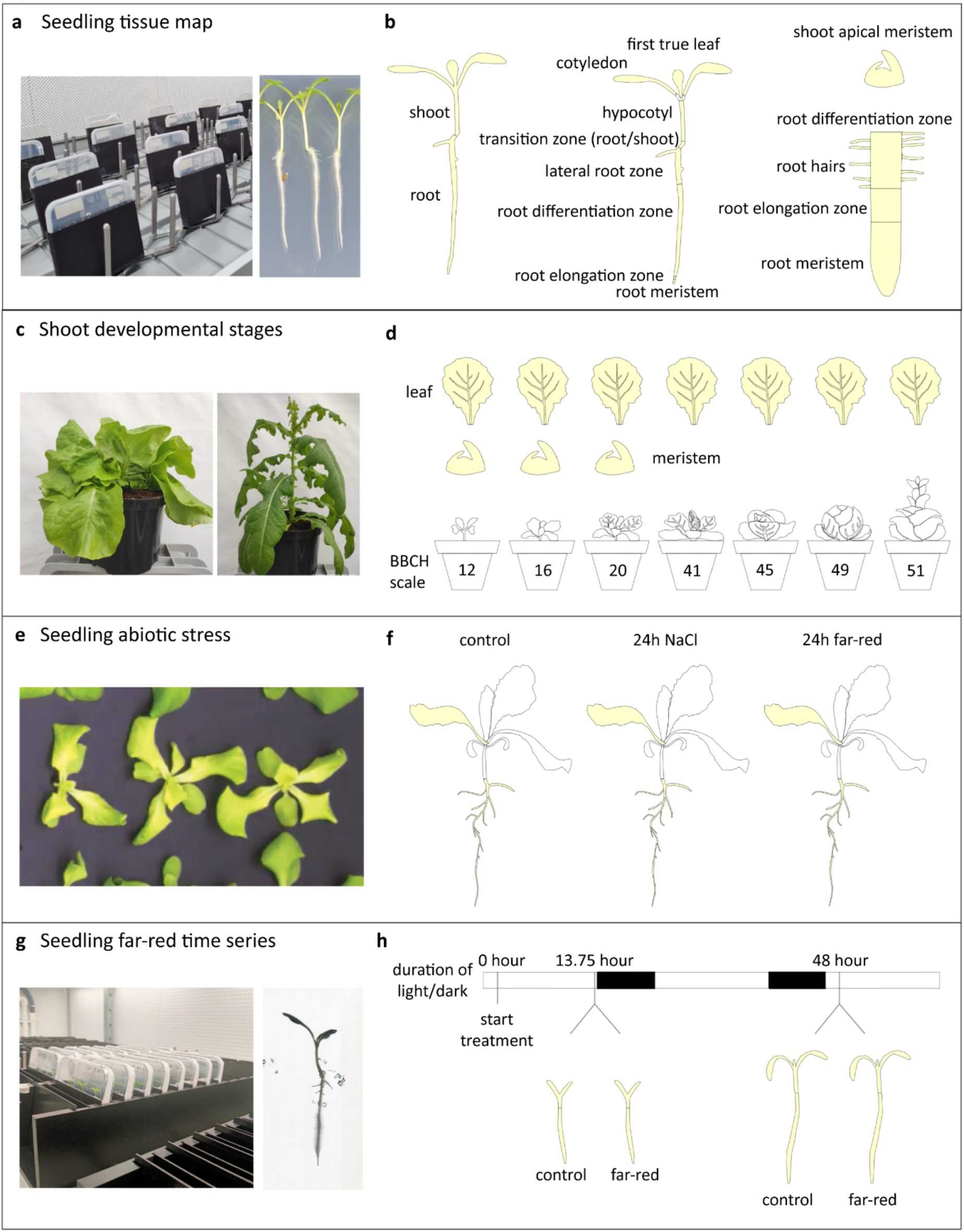
Overview of sample collection and experimental setup. **(a–b)** Seedling tissue map **(a)** Five *Lactuca* genotypes were grown on vertical plates with a dark-root system (left). 7-day-old *L. sativa* ‘Salinas’ seedlings are shown for reference (right). **(b)** Schematic representation of the twelve root and shoot tissues manually dissected from young seedlings of all genotypes. **(c–d)** Shoot developmental stages **(c)** *L. sativa* ‘Cobham Green’ at the folding stage (left) and *L. serriola* at the bolting stage (right), grown in pots under controlled conditions. **(d)** Sampling of young (1–2 cm) leaves and shoot apical meristems (SAM) at different developmental stages. SAMs were only collected from younger plants. **(e–f)** Seedling abiotic stress **(e)** Hydroponic growth setup for abiotic stress experiments. **(f)** Four *Lactuca* genotypes were grown under control conditions or exposed to two abiotic stress treatments - shade (far-red light) or salt (100 mM NaCl) - for 24 hours. Bulk root and shoot samples were collected at the end of the treatment. **(g–h)** Seedling far-red time series **(g)** *L. sativa* ‘Salinas’ seedlings were grown on vertical plates with a dark-root system (left) and exposed to either white light (control) or far-red-supplemented light. On the right, a ‘Salinas’ seedling is shown after 48 hours of far-red treatment. **(h)** Sampling time points for the time-series experiment. Root and shoot samples were collected at 13.75 hours (15 minutes before dusk) and at 48 hours (2 hours after dawn).

#### 1. Seedling tissue map

To achieve high spatial resolution of gene expression patterns, we finely dissected various root and shoot organs and tissues from young seedlings aged 7-11 days. Five different *Lactuca* genotypes were included: *L. sativa* ‘Salinas’, *L. sativa* ‘Cobham Green’, *L. serriola* US96UC23, *L. saligna* CGN05327, and *L. virosa* CGN04683. These seedlings were grown in a vertical plate system with a dark-root setup (**Figure 1a**). To account for differences in growth rates, *L. saligna* and *L. virosa* were grown for an extended period to match the seedling size of the other genotypes (**Figure S1**). Twelve distinct organs and tissues were manually dissected from all genotypes (**Figure 1b**): first true leaf, cotyledon, hypocotyl, transition zone, lateral root zone, root differentiation zone, root elongation zone, root and shoot meristems, root hairs, whole root and whole shoot.

#### 2. Shoot developmental stages

To expand our dataset and include developmental variation in gene expression, we sampled shoot samples from plants at different stages of growth, ranging from the two-leaf stage to bolting. As plants transition from early vegetative phases to later stages of development, their gene expression profiles shift. To capture this variation, we cultivated four *Lactuca* genotypes (*L. sativa* ‘Salinas’, *L. sativa* ‘Cobham Green’, *L. serriola* US96UC23, and *L. saligna* CGN05327) in soil-filled pots in a greenhouse until the bolting stage (**Figure 1c**). At specific developmental stages, defined by the BBCH scale for leaf vegetables forming heads (Meier 2018), we sampled young leaves (∼1–2 cm) and for the younger plants we additionally sampled tissue enriched for shoot apical meristems (SAMs; **Figure 1d**). Developmental stages were sampled as far as the architecture of the plants allowed (**Table S1**). For instance, heading stages were absent in *L. saligna* and *L. serriola*, as these species do not form heads.

#### 3. Seedling abiotic stress

We further extended our dataset by incorporating root and shoot samples from *L. sativa* ‘Olof’ seedlings grown hydroponically (**Figure 1e**). In addition to control conditions, two abiotic treatments were applied: supplemental far-red light exposure (simulating a low red to far-red light ratio; see **Figure S2** for light spectra) and salt stress, induced by adding 100 mM NaCl to the growth medium. To capture early transcriptional responses before significant developmental changes occurred, seedlings were exposed to the treatments for only 24 hours before bulk root and leaf tissues were collected (**Figure 1f**).

#### 4. Seedling far-red time series

Finally, we performed a time-series experiment with two time points to investigate the effects of supplemental far-red light exposure and time-of-day on gene expression in shoots and roots. *L. sativa* ‘Salinas’ seedlings were grown on vertical plates with a dark-root system (**Figure 1g**). Two-day-old seedlings were exposed to either white light (control) or white light supplemented with far-red light, and samples were collected at two time points: after 13.75 hours (15 minutes before dusk), and after 48 hours (2 hours after dawn) (**Figure 1h**). In addition to providing insights into the far-red light response, this dataset captures information on gene-expression patterns at two physiologically important times of day in lettuce seedlings.

### Overview of transcriptome data

In total, we obtained transcriptome data for 346 samples, including six genotypes, twelve organs/tissues and two treatments. Principal Component Analysis (PCA) of each dataset (**Figures 2a–e** for *L. sativa* and *L. serriola*, **Figure S3** for *L. saligna* and *L. virosa*) confirmed that biological replicates clustered closely together, reflecting high consistency in gene expression within conditions. The primary drivers of variation were organ type, developmental stage or genotype. PCA of the “Seedling tissue map” dataset showed that samples clustered logically with similar tissues grouping closely together across *L. sativa* and *L. serriola* accessions (**Figure 2a**). This indicates conserved organ development, while additional cultivar- or species-specific clustering within each tissue cluster suggests subtle transcriptomic differences between genotypes. Similarly, in the “Shoot developmental stages” dataset, PCA revealed clear separation between the two tissue types (**Figure 2b**). PC1, which explains 47% of the variation, and PC2 (14.5%) capture tissue identity (leaf vs. SAM) and developmental progression. The “Seedling abiotic stress” dataset also showed clear separation of samples by tissue type and treatment (**Figure 2c**). PC1, which explains 76.6% of the variation, reflects organ identity (leaf vs. root), while PC2 (4.6%) represents the treatment effect. The impact of salt stress seemed more pronounced than that of far-red supplemental light in both leaves and roots. In the “Seedling far-red time series” dataset, PC1 (66.5%) again reflects organ identity (leaf vs. root; **Figure 2d**), while PC2 (9.4%) captures variation mainly associated with time-of-day rather than the far-red treatment (**Figure 2d-e**). Separating the root and leaf samples, batch and time-of-day effects are visible in the first two principal components (**Figure S4**).

**Figure 2:**
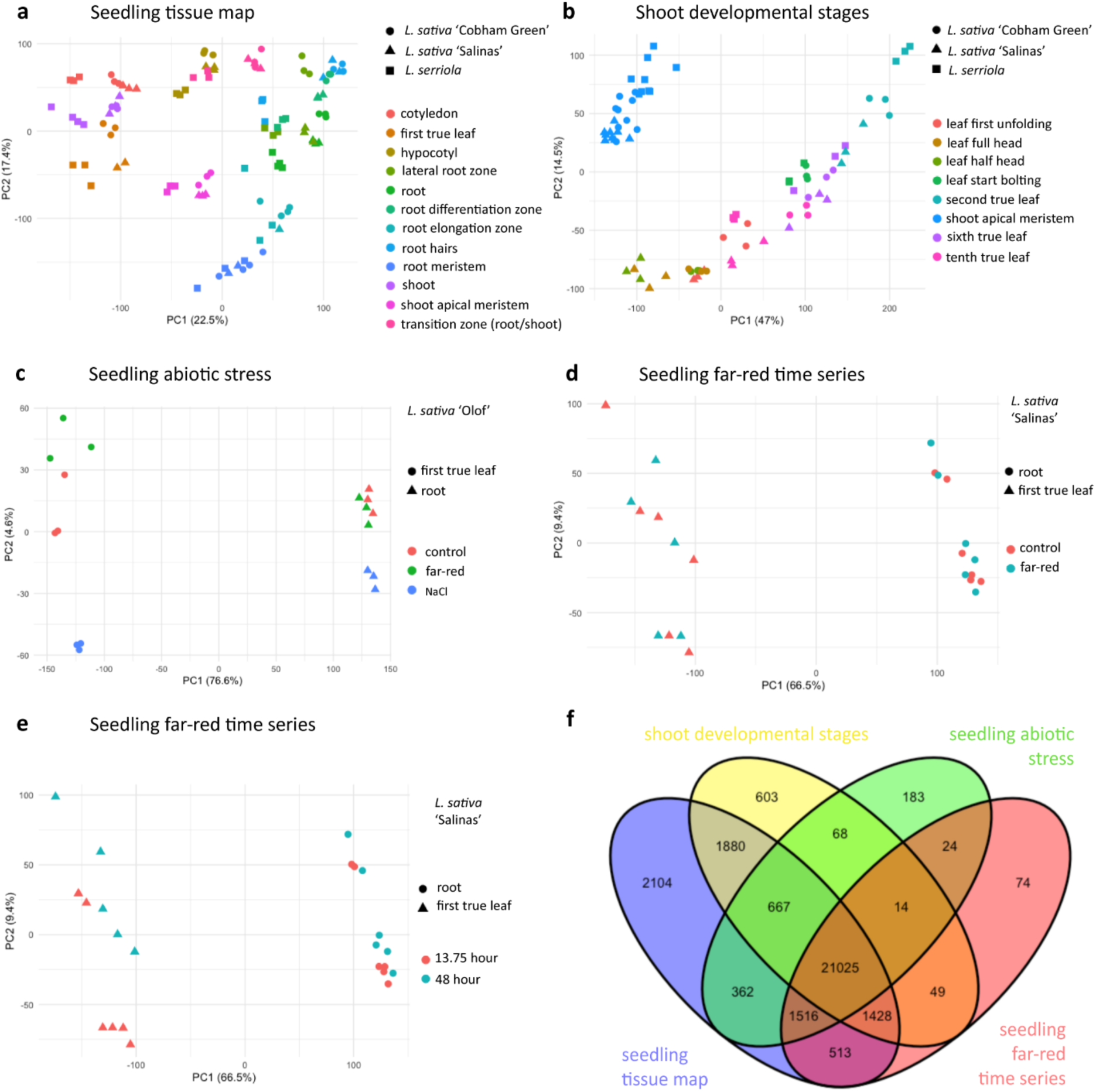
Global overview of gene expression patterns across *L. sativa* and *L. serriola* datasets. **(a-d)** Principal component analysis (PCA) plots showing the variation in gene expression across samples within each experiment: **(a)** Seedling tissue map, **(b)** Shoot developmental stages, **(c)** Seedling abiotic stress, and **(d-e)** Seedling far-red time series. **(f)** Venn diagram depicting the number of genes expressed (TPM ≥ 2 in at least two samples) in each dataset and their overlap across experiments.

Gene expression levels were assessed across all samples, with *L. sativa* and *L. serriola* directly comparable due to mapping against the same *L. sativa* reference genome. In total, 30,510 genes had a TPM (transcripts per million) ≥ 2 in at least two samples, representing about 60% of the annotated genes in *L. sativa* ‘Salinas’ (**Figure 2f**). PCA plots and numbers of expressed genes (TPM ≥ 2) for *L. saligna* and *L. virosa* samples, which were included in selected experiments, are provided in **Figure S3**. The “Seedling tissue map” experiment had the highest number of expressed genes (29,495), likely due to the inclusion of multiple genotypes and a diverse range of organs and tissues. In contrast, the “Seedling abiotic stress” experiment had the lowest number of expressed genes (23,859). This is consistent with the use of a single genotype and very young seedlings with limited organ differentiation. The “Shoot developmental stages” and “Seedling far-red time series” experiments had 25,734 and 24,643 expressed genes, respectively.

A core set of 21,025 genes was expressed across all four datasets for the *L. sativa* and *L. serriola* genotypes, while each dataset also contained uniquely expressed genes. Notably, the “Seedling tissue map” experiment had the highest number of dataset-specific genes, with 2,104 genes expressed exclusively in this dataset. This highlights the value of fine tissue dissection for capturing localised gene expression. For instance, expression of a homolog of *LOB DOMAIN-CONTAINING PROTEIN 1* (*LBD1*, LOC111881535) was detected in the shoot apical meristem of only this experiment (**Figure S5a**). Likewise, a homolog of *HIGH AFFINITY NITRATE TRANSPORTER 2.6* (*NRT2.6*, LOC111885982) was expressed most strongly in the root elongation zone, illustrating highly localised expression patterns **(Figure S5b**). Expression of another *NRT2.6* homolog (LOC111886060) showed preferential expression in the root elongation zone of *L. serriola* (**Figure S5c**), underlining species-specific differentiation in gene expression.

### Design and features of the Lettuce Expression Browser

The LEB was designed to host multiple datasets, and to visualise and compare gene expression levels across diverse lettuce datasets. Since each experiment was conducted independently, the data are organised into separate tabs to emphasise their distinct experimental contexts (**Figure 3a-b**). Illustrations used in the visualisations were created from photographs taken during the experiments, ensuring accurate representation of plant morphology at each sampling point. These images depict control plants at the time of sampling; treatment-induced phenotypic changes are not shown.

**Figure 3:**
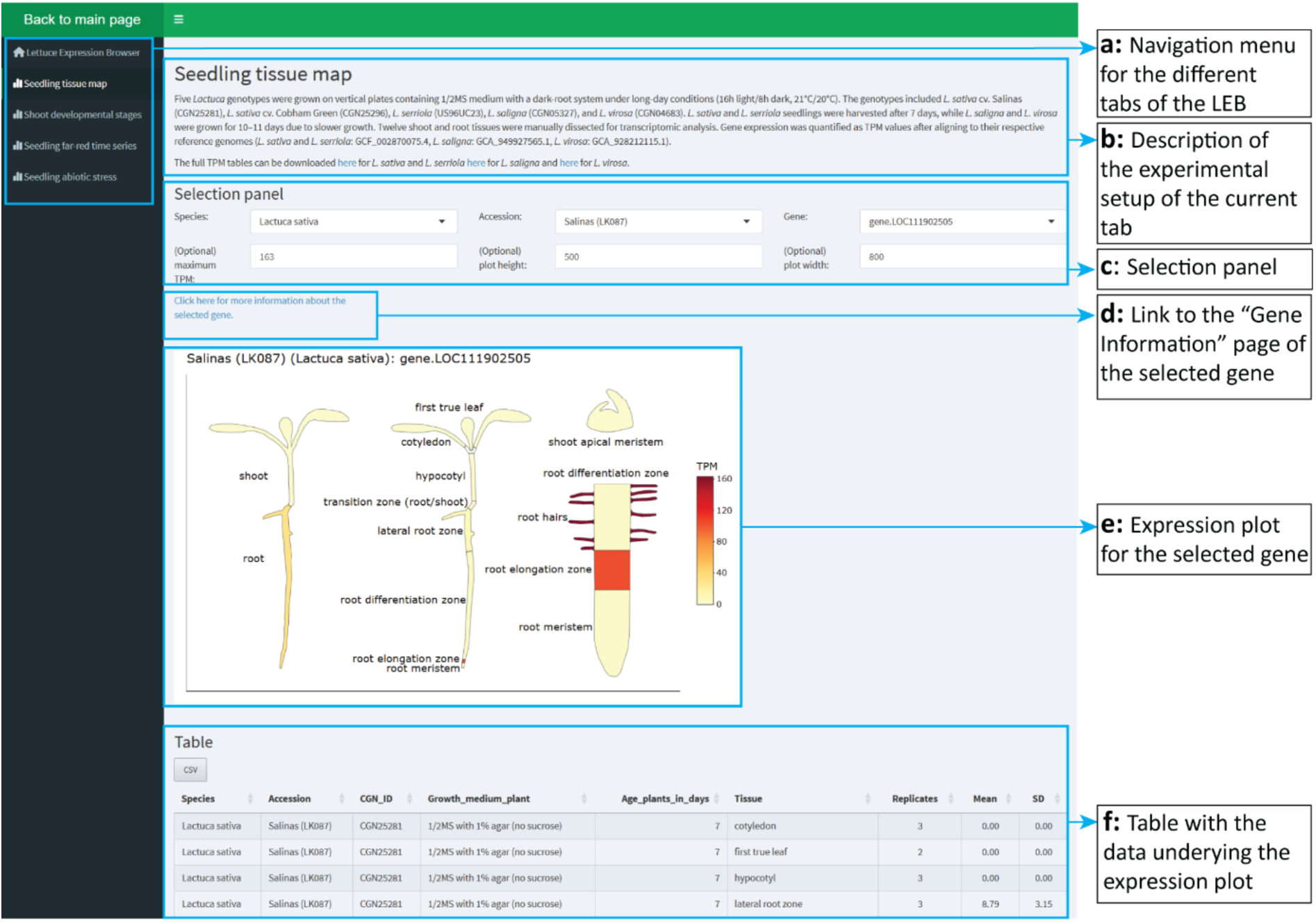
Lettuce Expression Browser (LEB) interface overview. Screenshot of the LEB showing the expression pattern of LOC111902505 in *L. sativa* ‘Salinas’ in the “Seedling tissue map”. Functional sections of the interface **(a-f)** are highlighted, with annotations explaining the purpose and features of each component.

Averaged gene expression levels (across replicates) were visualised using a white–yellow–red colour scale (**Figure 3e)**, a common convention in gene expression studies (Waese et al. 2017), and quantified in TPM to reflect transcript abundances (Li and Dewey 2011). This allows direct comparison of expression levels across both samples and genes. To support interpretation of the expression data, we included standard deviation values and the number of underlying replicates for each data point in the visualisation. The LEB interface allows users to adjust display parameters (e.g. colour scale and plot size; **Figure 3c**) and to download both figures and corresponding data tables (**Figure 3f**). Developed in close collaboration with experimental researchers, the LEB underwent extensive user testing and refinement (**Methods S1**). It was found to be informative and user-friendly by the both the original researchers and independent users unfamiliar with the experiments.

Common identifiers are crucial for effective querying within the LEB. For gene identifiers, we used the *L. sativa* ‘Salinas’ v11 RefSeq annotation, supplemented with non-overlapping genes from the GenBank annotation of the same genome assembly (van Workum, de Ridder, et al. 2024). At the time of LEB construction, no other publicly available and well-annotated genome assemblies existed for the *L. sativa* and *L. serriola* accessions used to generate the LEB other than *L. sativa* ‘Salinas’, which is why it served as the reference for both species. For *L. saligna* and *L. virosa*, gene identifiers were derived from their respective GenBank structural annotations (Xiong, van Workum, et al. 2023; Xiong, Berke, et al. 2023).

To support the interpretation of gene expression across tissues and conditions, functional descriptions of genes are essential. Therefore, we developed a “Gene Information” page (**Figure 3d**) that enables querying of genes and functional domains in all *Lactuca* spp. reference genomes as well as *Arabidopsis thaliana* (ARAPORT11). The page provides both predicted functional annotations and homology information for each gene, allowing for an initial overview of gene characteristics and supporting translational analysis across species.

### Exploring gene expression patterns in lettuce

The browser is a powerful tool for visualising gene expression patterns across different tissues, treatments and developmental stages, allowing researchers to dissect processes with high resolution. By presenting spatially resolved expression data, it facilitates the identification of key regulatory genes and their specific roles in plant development.

For example, our data of the “Seedling tissue map” demonstrate that the lettuce homolog of *ROOT HAIR DEFECTIVE 6* (*RHD6*, LOC111902419), a key regulator of root hair specification, was specifically expressed in the root meristem and elongation zone (**Figure 4a**). This pattern closely mirrors that of its *Arabidopsis thaliana* counterpart (Menand et al. 2007), suggesting a conserved function in initiating root hair development in lettuce. Similarly, the lettuce homolog of *EXPANSIN A7* (*EXPA7*, LOC111902505), essential for root hair elongation in *Arabidopsis* (Lin, Choi, and Cho 2011), was predominantly expressed in the root elongation zone and isolated root hairs (**Figure 4a**). Interestingly, while *EXPA7* was highly expressed in isolated root hairs, its expression in the differentiation zone was minimal despite the presence of root hairs in this region, demonstrating how tissue-specific sampling can uncover precise expression domains that would otherwise be obscured in bulk organ analyses.

**Figure 4:**
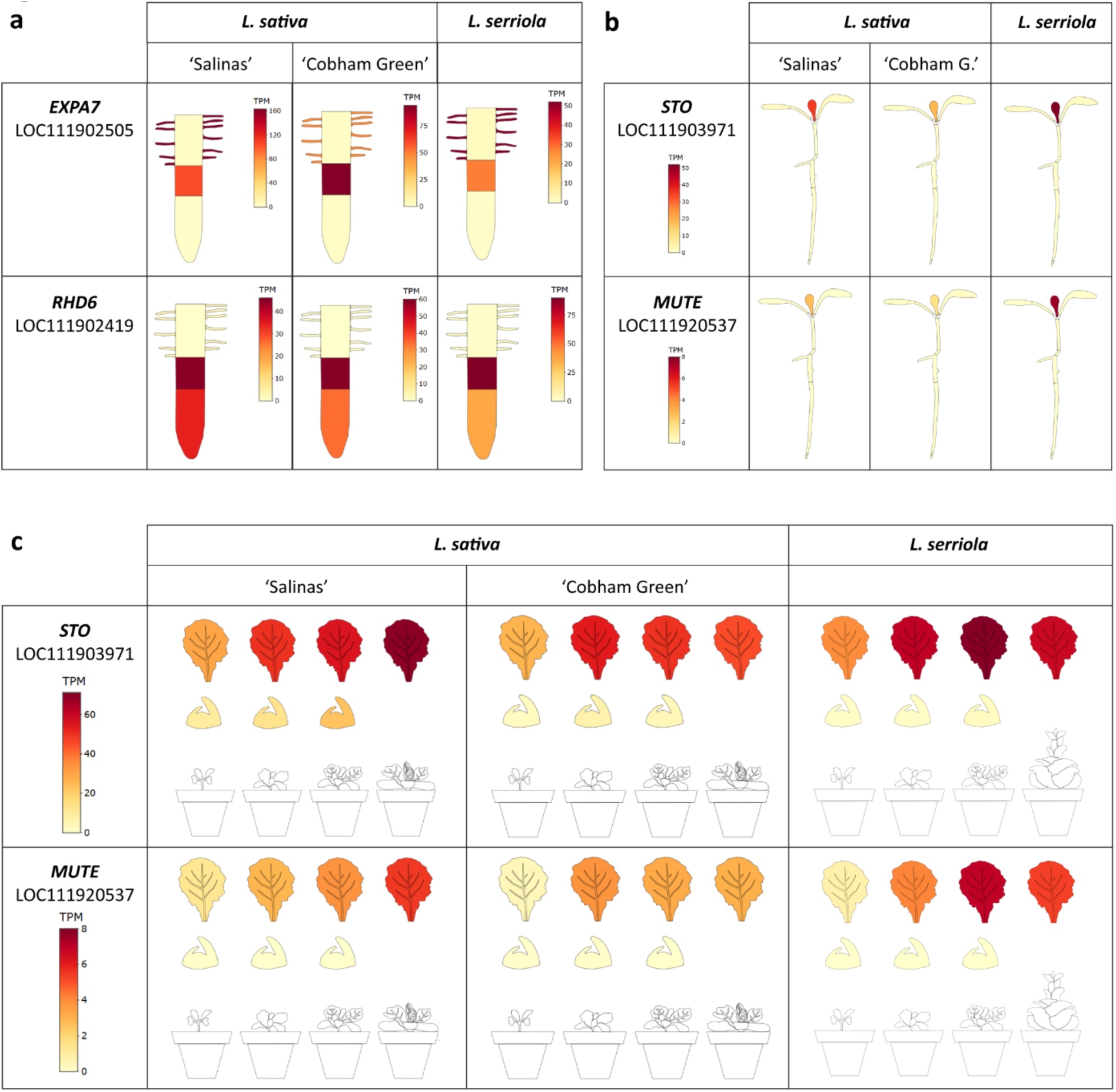
Spatial and temporal gene expression patterns in lettuce visualised using the expression browser. **(a)** Expression profiles of *RHD6* (LOC111902419) and *EXPA7* (LOC111902505) in root tissues reveal zone-specific expression patterns. *RHD6* is expressed in the meristem and elongation zone, while *EXPA7* shows highest expression in the elongation zone and isolated root hairs, consistent with their roles in root hair specification and outgrowth. **(b)** Organ-specific expression of *STOMAGEN* (*STO*) (LOC111903971) and *MUTE* (LOC111920537) in the “Seedling tissue map” dataset, showing specific expression in the first leaf. **(c)** Expression pattern of *STO* across the “Shoot developmental stages” dataset in different *L. sativa* cultivars and *L. serriola*, highlighting increased expression in later-developing leaves. TPM: transcripts per million.

The browser also facilitates the exploration of gene expression across organs and developmental stages, allowing researchers to investigate gene activity in different experimental contexts. For instance, analysis of the “Seedling tissue map” dataset revealed that orthologs of *STOMAGEN* (*STO*, LOC111903971) and *MUTE* (LOC111920537), key regulators of stomatal development, were specifically expressed in the first leaf within this dataset (**Figure 4b**). In Arabidopsis, *STO* is expressed in leaf mesophyll cells, where it functions as a secreted peptide promoting stomatal formation by antagonizing negative regulators, while *MUTE* expression is restricted to meristemoid cells, where it controls the transition from meristemoids to guard mother cells (Pillitteri et al. 2007; Sugano et al. 2010). Further investigation using the “Shoot developmental stages” dataset showed that both genes were also expressed in leaves formed at later growth stages (**Figure 4c**). Notably, in both *L. sativa* cultivars and the wild relative *L. serriola*, expression levels of *STO* and *MUTE* increased in later-developing leaves. *STO* and *MUTE* expression levels can correlate with stomatal density, as overexpression of *STO* increases stomatal numbers and knock-outs of either gene reduce them (Pillitteri et al. 2007; Sugano et al. 2010). Thus, the increased expression in leaves formed at later developmental stages suggested that later-formed leaves contain more stomata. Indeed, our analysis of stomatal density in *L. sativa* ‘Salinas’ and ‘Cobham Green’ confirmed that later-developing leaves exhibit a higher stomatal density compared to earlier-formed leaves (**Figure S6**). These findings suggest that stomatal density can serve as a marker for the juvenile-to-adult transition in lettuce, as observed in other species (Feng et al. 2016; Lawrence et al. 2021).

The “Seedling far-red time series” dataset provides insights into the transcriptional response to far-red light and its interaction with circadian regulation. The sampling strategy was designed to capture both the effect of far-red light on genes related to circadian rhythm regulation and the later transcriptional response to far-red light. Far-red light is perceived by phytochromes such as PHYTOCHROME B (phyB), which contributes to circadian clock regulation, and accordingly, *phyB* mutants displayed impaired clock-regulated responses (Romanowski et al. 2021; Casal and Yanovsky 2005). In accordance, we observed higher expression of two of the six *L. sativa* orthologs of the *Arabidopsis* clock component *GIGANTEA* (*GI;* LOC111891398 and LOC111919525) upon far-red treatment (**Figure 5a**).

**Figure 5:**
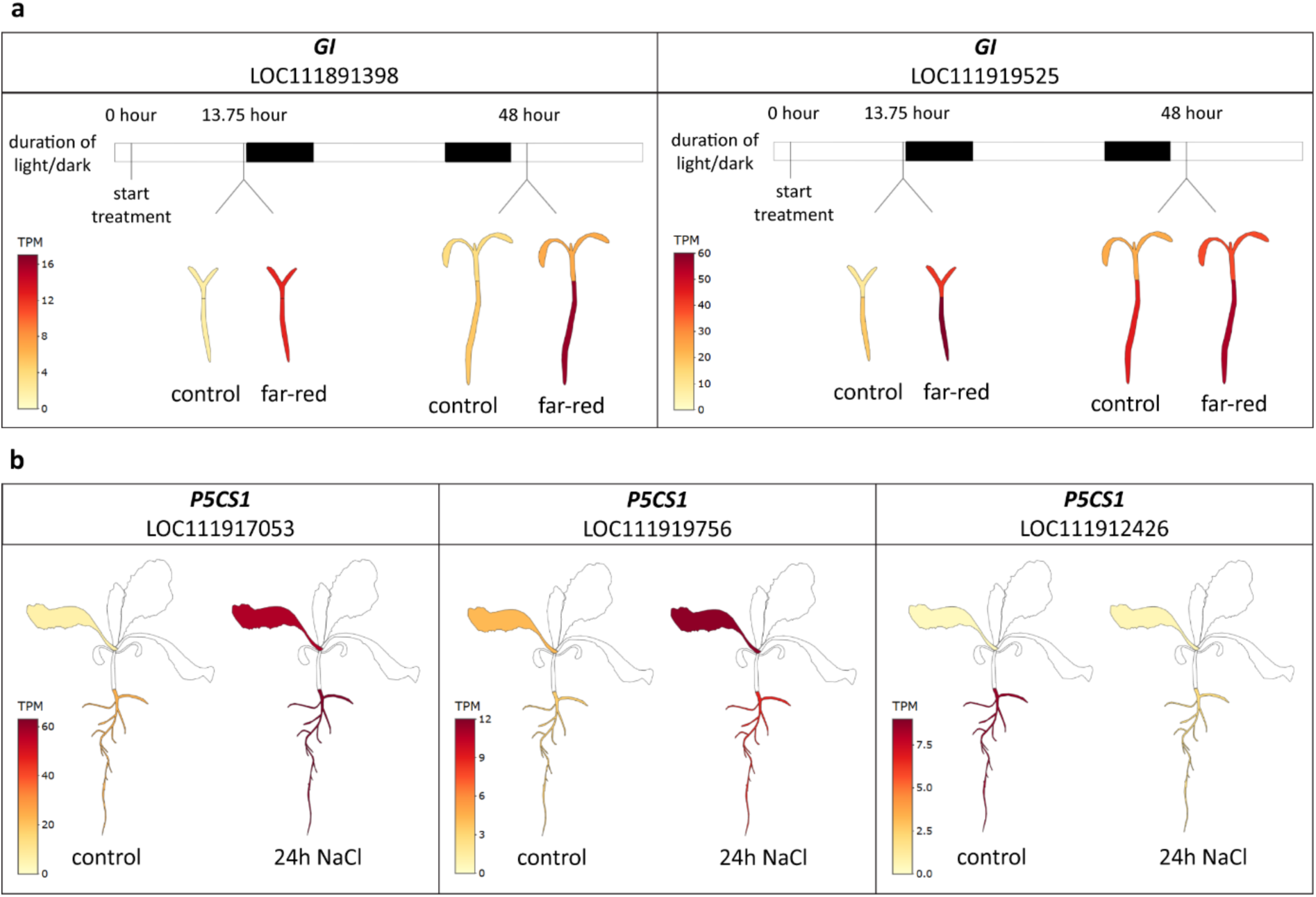
Effects of far-red light and salt treatment on gene expression in lettuce. **(a)** Temporal expression of two lettuce *GIGANTEA* (*GI*) orthologs (LOC111891398 and LOC111919525) in the “Seedling far-red time series” dataset, showing elevated expression in *L. sativa* ‘Salinas’ shoots and roots after 13.75 hours of far-red light treatment. **(b)** Expression patterns of three lettuce orthologs of *P5CS1* in roots under control and salt stress conditions. Expression data are from the “Seedling abiotic stress” dataset, showing transcript abundance in roots of *L. sativa* ‘Olof’ after 24 hours of salt treatment or control conditions. Two homologs (LOC111917053 and LOC111919756) are strongly upregulated in response to salt stress, whereas the third homolog (LOC111912426) is downregulated. TPM: transcripts per million.

The “Seedling abiotic stress” dataset contains expression profiles from both roots and the first leaf under 24-hour treatments with salt and supplemental far-red light. Focussing specifically on the salt treatment, we examined the expression of three lettuce homologs of *DELTA1-PYRROLINE-5-CARBOXYLATE SYNTHASE 1* (*P5CS1*), a central gene in proline biosynthesis and salt stress response in *Arabidopsis* (Székely et al. 2008). Two of the three homologs (LOC111917053 and LOC111919756) are upregulated in roots and first leaf after 24 hours of salt exposure, whereas they show little to no expression under control conditions (**Figure 5b**). In contrast, a third homolog (LOC111912426) is specifically expressed in roots under control conditions and is downregulated upon salt treatment (**Figure 5b**). These contrasting patterns illustrate how the browser enables exploration of differential regulation among homologs, supporting functional predictions and suggesting potential subfunctionalisation within gene families.

### Cross-species gene expression analysis

The LEB supports cross-species comparisons of gene expression, helping to identify conserved or divergent patterns across *Lactuca* species. As an example, we used the tool to explore expression of *DEEPER ROOTING 1* (*DRO1*), a gene first identified in rice where it regulates root growth angle and improves drought tolerance when overexpressed (Uga et al. 2013; Uga, Okuno, and Yano 2011). Homologues in *Arabidopsis thaliana* (*DRO1/LZY3* and *DRO3/LZY2*) have similar roles in root and shoot gravitropism (Taniguchi et al. 2017; Guseman et al. 2017; Yoshihara and Spalding 2017; Ge and Chen 2016).

Homologues of *A. thaliana DRO1* were identified in lettuce using the Gene Information page, and their expression profiles were visualised across species and cultivars (**Figure 6**). In *L. sativa*, *L. serriola*, and *L. virosa*, two homologues were found. Unlike *Arabidopsis*, where both *DRO1/LZY3* and *DRO3/LZY2* are expressed in root and hypocotyl (Taniguchi et al. 2017), lettuce homologues show clear expression divergence: LOC111882727 is predominantly expressed in the root tip, while LOC111921379 is expressed in both root and hypocotyl **(Figure 6a)**. In *L. sativa* cultivars (‘Salinas’ and ‘Cobham Green’), LOC111921379 showed relatively higher expression in the hypocotyl than in the root tip. In *L. serriola* and *L. virosa*, expression of this homologue was more balanced between the two tissues. In *L. saligna*, only a single *DRO1/3* homologue was identified, with expression in both root and hypocotyl, suggesting a possible loss of the root-specific copy or gaps in the genome assembly. A third related gene, LOC111892617, corresponding to *Arabidopsis DRO2/LZY4*, was also identified and found to be consistently root-specific across all *Lactuca* species examined (**Figure 6b**), similar to its expression pattern in Arabidopsis (Yoshihara and Spalding 2017). These patterns demonstrate how the LEB can be used to uncover expression divergence and infer evolutionary changes in gene regulation and conserved functional domains across species.

**Figure 6:**
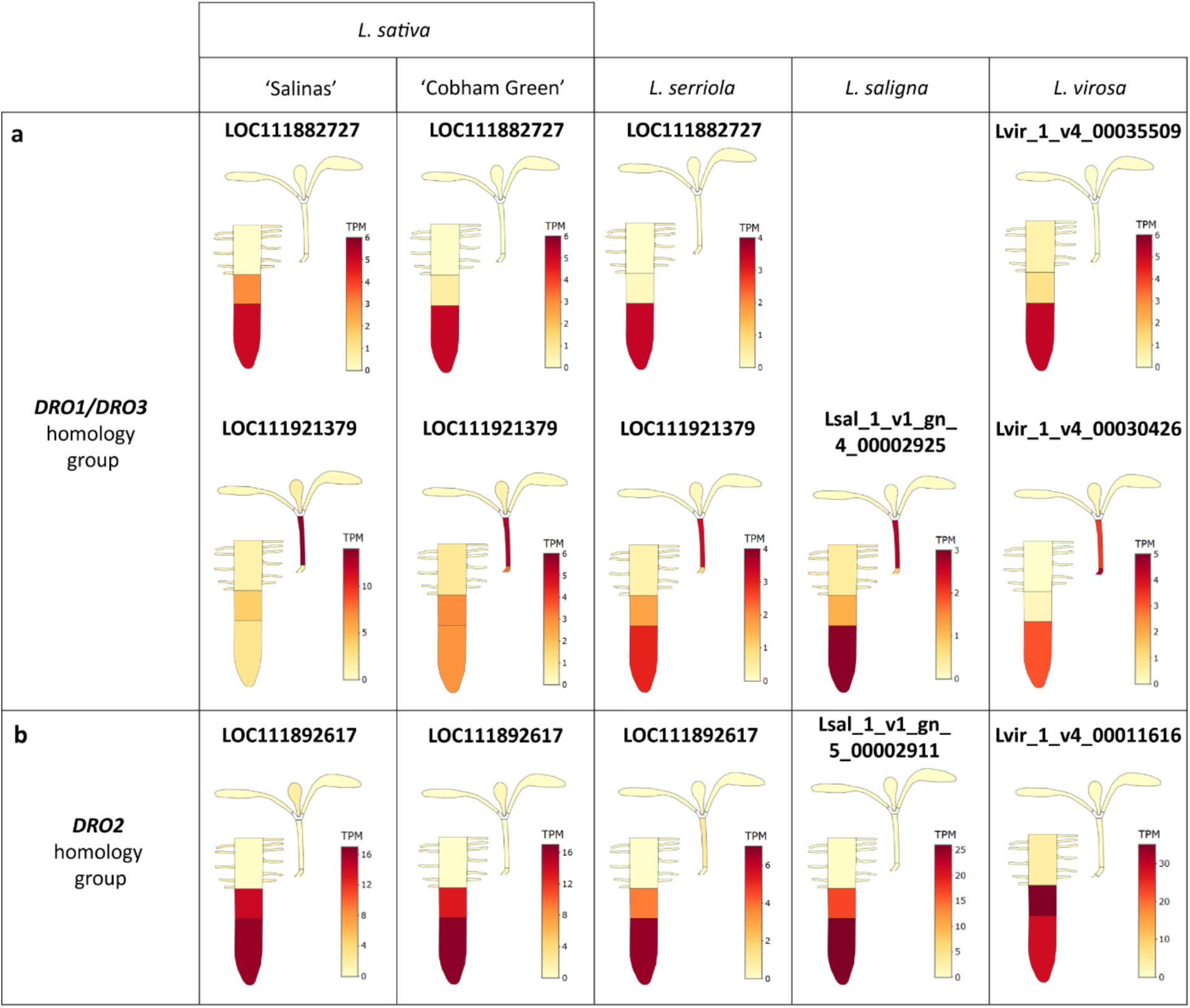
Cross-species analysis of *DRO1* homologue expression in *Lactuca* species. **(a–b)** Gene expression patterns of *DRO1* homologues in *Lactuca* species from the “Seedling tissue map” dataset. Expression levels are shown for the root tip and the shoot in *L. sativa*, *L. serriola*, *L. saligna*, and *L. virosa*. **(a)** Two homologues of *Arabidopsis DRO1/DRO3* are present in all species except *L. saligna*, which contains only one. One homologue shows root-specific expression, while the other is expressed in both root tip and hypocotyl. In *L. sativa* cultivars, the latter shows relatively higher expression in the hypocotyl, while in *L. serriola* and *L. virosa*, expression of this homologue is more balanced between the two tissues. **(b)** A third homologue, corresponding to *Arabidopsis DRO2*, is consistently root-specific across all species.

## Discussion

We introduce the Lettuce Expression Browser (LEB), a comprehensive, modular platform for exploring gene expression patterns in *Lactuca* species. Despite its growing importance as a model species in the Asteraceae family, lettuce lacks the high-resolution, context-rich transcriptomic resources available for other crops. Existing platforms are limited in scope, often based on a single cultivar and lacking the tissue-specific and condition-dependent detail necessary to support functional genomics and breeding applications (Zhang et al. 2017; Guo et al. 2023). Gene expression is highly dynamic: shaped by genotype, developmental stage and environmental stimuli. The ability to interrogate expression patterns in these specific contexts allows for the discovery of key regulatory genes and pathways. The LEB addresses this gap by offering a detailed and interactive expression atlas that enables comparative, condition-aware exploration of gene activity. It enables researchers to explore when, where, and in which genotype, a gene is expressed, offering critical insights into its potential biological role.

The current version of the LEB includes four independent experimental setups, primarily based on the cultivar ‘Salinas’, which serves as the reference genome (Reyes-Chin-Wo et al. 2017). However, most datasets also incorporate additional genotypes, allowing for comparative analyses. While the seedling stage is a common focus, one dataset extends beyond this phase, demonstrating the LEB’s potential for broad developmental coverage. The browser’s interface draws on established expression browsers from other species (Waese et al. 2017), supporting usability and accessibility for researchers across disciplines.

A key feature of the LEB is its modular architecture, which allows for integration of future datasets. This makes the platform scalable and adaptable, enabling it to grow alongside ongoing transcriptomic efforts. In particular, the inclusion of data from biotic stress experiments will be important for uncovering regulatory pathways relevant to disease resistance and other agriculturally significant traits. However, despite the modular setup of the LEB, currently no method exists for external users to upload publicly available data. Automated creation of experimental visualisations cannot feasibly be done in an accurate manner, because it requires manual steps to connect expression data with the expression maps based on metadata.

Understanding the differences in gene expression across cultivars and even between species is crucial for linking genes to their functions in specific biological contexts. The LEB facilitates this by providing access to homologous gene information through its integrated Gene Information feature. This allows for exploration of differences in gene expression between accessions. However, quantifying reads of one accession against a reference genome of another accession still creates biases in expression levels (Zhan, Griswold, and Lukens 2021). Pangenome-based comparisons using pangenome mapping (Sibbesen et al. 2023) or comparisons using homology are possible approaches (Garassino et al. 2024). On the other hand, resources such as the Reference Transcriptome Datasets (RTD) that unify transcript diversity into one compiled set of transcripts (Kara et al. 2024) are important steps in the direction of obtaining complete transcriptome datasets, but lack expression diversity across accessions. While the LEB currently relies on gene-level quantification based on the ‘Salinas’ reference genome, future updates may include transcript-level views, multi-genotype alignment strategies or pantranscriptomic approaches. Advances like these are key to unlocking the full potential of lettuce transcriptomics and applying it to breeding innovation.

## Materials and methods

### Plant material, growth conditions and sample collection

#### Seedling tissue map

The *Lactuca* lines used in this experiment included varieties *L. sativa* ‘Salinas’ (CGN25281) and *L. sativa* ‘Cobham Green’ (CGN25296), and wild-collected accessions of *L. serriola* (US96UC23), *L. saligna* (CGN05327) and *L. virosa* (CGN04683).

Seeds were surface sterilised by immersing them in 50% household bleach for 5 minutes, followed by five rinses with sterile MQ water. Ten seeds were then placed on 12 cm square plates containing ½MS: half-strength Murashige and Skoog medium including vitamins (Duchefa Biochemie) with MES buffer and 1% agar, pH 5.8, without sucrose. Plates were kept at 4°C for 6-7 days for stratification. The plates were positioned vertically and single-plate cardboard covers were used to ensure a dark-root system. Seedlings were grown under long-day conditions (16-hour light/8-hour dark) at 21°C during the day and 20°C at night, with a light intensity of ∼150 µmol/m²/sec and 70% relative humidity. *L. sativa* and *L. serriola* seedlings were harvested after 7 days, while *L. saligna* and *L. virosa* seedlings were grown for 10 and 11 days, respectively, to account for their slower growth rates (**Figure S1**).

All tissues and replicates of a given genotype were sampled on the same day with three replicates per sample type, with a few exceptions (**Table S2**). Sample collection was performed in the early afternoon. Seedling tissues were manually dissected using tweezers, scalpels and binoculars, and pooled as indicated (**Table S3).** Most tissues were immediately snap-frozen in liquid nitrogen (N_2_) and stored at - 80°C. Tissue types that required extensive sampling were first collected in 200 µL RNAlater™ Stabilization Solution (Invitrogen) at room temperature, followed by removal of the RNAlater before snap-freezing. (**Table S3**).

Root hairs were isolated as described previously (Bucher et al. 1997), with modifications. Root systems (excluding the meristematic region) were stirred with a plastic rod in a clean ice bucket filled with N_2_. After ∼20 minutes, when root hairs had detached and floated in the N_2_, the suspension was filtered through a course metal sieve (∼1 mm pore size) multiple times to remove large root fragments. Smaller root debris was removed by subsequent sieving through a finer mesh (∼0.5 mm pore size). The remaining root hair-enriched fraction was transferred into a 50-ml tube, and after N_2_ evaporation, the root hairs were stored at −80°C.

#### Shoot developmental stages

The *Lactuca* lines used in this experiment included *L. sativa* ‘Salinas’ (CGN25281), *L. sativa* ‘Cobham Green’ (CGN25296), *L. serriola* (US96UC23) and *L. saligna* (CGN05327). Plants were grown in a greenhouse at the Delphy Improvement Centre (Greenhouse Horticulture, Bleiswijk, The Netherlands) under semi-controlled conditions. While key environmental parameters were regulated, greenhouse conditions inherently introduced variability, including minor fluctuations in temperature, humidity and light quality. Seedlings were germinated on peat quickplugs and grown in 5L pots filled with potting mix. The growth conditions were as follows: long-day photoperiod (16 hours light / 8 hours dark), day temperature of 23°C, night temperature of 17°C, 60% humidity, and a photosynthetic photon flux of 12 mol/m²/day. During the day, CO_2_ was supplemented to a concentration of 800 ppm with no CO_2_ supplementation at night.

Plant development was assessed using the BBCH scale, which describes the phenological stages of leafy vegetables forming heads (Meier 2018). The stages evaluated in this study included BBCH stage 12 (2 leaves), BBCH stage 14 (4 leaves), BBCH stage 16 (6 leaves), BBCH stage 20 (10 leaves), BBCH stage 41 (2 leaves are folding), BBCH stage 45 (half head size), BBCH stage 49 (full head size) and BBCH stage 51 (bolting, characterised by the onset of stem elongation). The time required to reach each developmental stage differed between genotypes (**Table S1**).

Samples were collected from three replicates per genotype on the same day. For each developmental stage, young leaves (∼1-2 cm) were sampled and a 5-mm diameter disc was excised from each leaf. Leaf discs from four plants were pooled to create a single sample. Shoot apical meristems (SAMs) were excised from the younger stages using binoculars but due to the technical limitations of manual dissection, these samples should be considered meristem-enriched rather than exclusively meristematic. Four SAMs were pooled per sample. All samples were snap-frozen in a slurry of 96% ethanol and dry ice and transported on dry ice. After collection, samples were stored at −80°C for further processing. Notably, plants of *L. sativa* ‘Cobham Green’ and *L. saligna* at the latest developmental stages exhibited signs of aphid infestation and downy mildew infection, which may have influenced gene expression profiles. These samples are marked with a disclaimer in the LEB to ensure transparency for users analysing the dataset.

### Stomatal density analysis

In addition to gene expression profiling, stomatal density was quantified in *L. sativa* cultivars ‘Salinas’ and ‘Cobham Green’ grown under controlled environmental conditions. Plants were cultivated in climate-controlled growth chambers under a long-day photoperiod (16 h light / 8 h dark) using BX180c3 white LED light at 200 µmol m⁻² s⁻¹ (Valoya, Helsinki, Finland), 20°C, and 70% relative humidity. Seeds were stratified for three days and transplanted to pots containing soil, which were covered with transparent plastic sheets for the first seven days after transplanting.

At 35 days after germination (DAG), the first up to the tenth leaves were detached and leaf dimensions were recorded. To obtain imprints, a thin layer of clear nail polish was applied between the midrib and the blade edge on both the adaxial and abaxial surfaces of fully expanded leaves. After drying, the nail polish layer was carefully removed using clear adhesive tape and mounted on microscope slides. Imprints were examined using a Zeiss Axioscope A1 compound microscope (Zeiss, Oberkochen, Germany) under phase contrast at 10× and 20× magnification. Stomata were counted manually from two independent imprints per leaf, and the values were averaged. Mean stomatal density (stomata mm⁻²) was calculated for each leaf position and genotype (n > 10) using FIJI software (Schindelin et al. 2012).

#### Seedling abiotic stress

*Lactuca sativa* ‘Olof’ (CGN05786) were grown hydroponically in 1× Hyponex medium under long-day conditions (16-hour light/8-hour dark) at 21°C during the day and 18°C at night, with a light intensity of ∼200 µmol/m²/sec (PAR). 14-day-old seedlings were treated for 24 hours. Seedlings were subjected to one of three treatments: a control condition in which seedlings continued to grow under the initial conditions, a salt stress condition in which 100 mM NaCl was added to the hydroponic solution, and a far-red light condition in which far-red light was added to the standard light spectrum, increasing the total photon flux to ∼220 µmol/m²/sec (PAR) with a R:FR ratio of 0.15 (**Figure S2a-b**). After 24 hours of treatment, plant samples were harvested. For shoot samples, 6-mm leaf discs were collected per plant and pooled from three plants. Root samples were collected and pooled from the same three plants. All samples were snap-frozen in liquid nitrogen and stored at −80°C for further processing.

#### Seedling far-red time series

For this experiment, *L. sativa* ‘Salinas’ (CGN25281) seeds were first stratified at 4°C for 7 days. The seeds were then surface-sterilised and plated on ½MS medium as described above. The plates were positioned vertically in a rack that covered the roots to ensure a dark-root system (Figure 1g). Growth conditions were set to a long-day cycle with temperatures of 21°C during the day and 20°C at night, a 16-hour light period (∼150 µmol/m²/sec) and 70% humidity. Two days post-germination, the seedlings were transferred to control conditions (White light: Valoya LightDNA (NS1/FR), PAR: ∼150 µmol/m^2^/sec) or far-red supplemented light (Valoya LightDNA (FR), PAR: ∼150 µmol/m^2^/sec, R:FR=0.3) (https://www.valoya.com/lightdna/) (**Figure S2c-d**).

Root and shoot samples were collected after 13.75 and 48 hours of treatment, with each sample consisting of four pooled seedlings. Samples were immediately snap-frozen in liquid nitrogen for downstream analysis.

### RNA extraction and sequencing

For each sample, total RNA was extracted using the MagMAX™ Plant RNA Isolation Kit (2% (w/v) PVP40 added to the Lysis Buffer) on a Kingfisher robot. RNA sequencing was performed at Utrecht Sequencing Facility (USEQ) on a NovaSeq6000 for all experiments (except for the “Seedling abiotic stress” samples which were run on a NextSeq 2000). Libraries were enriched using Illumina TruSeq polyA, stranded and paired-end (100bp).

### Processing of RNA-seq data

The current v11 *L. sativa* ‘Salinas’ reference genome (Reyes-Chin-Wo et al. 2017) has two annotations available: the original Genbank annotation (GCA_002870075.4) and a RefSeq annotation (GCF_002870075.4). We used a merged annotation to capture all potentially expressed genes (van Workum, de Ridder, et al. 2024). For *L. saligna* and *L. virosa* samples, we used their respective reference genomes from Genbank without modification: GCA_949927565.1 (Xiong, Berke, et al. 2023) for *L. saligna* and GCA_928212115.1 (Xiong, van Workum, et al. 2023) for *L. virosa*. *L. serriola* reads were mapped against the *L. sativa* genome, as no *L. serriola* genome was available. RNA-seq reads were processed with the RNASeq-NF pipeline at Utrecht Bioinformatics Expertise Core (UBEC) (https://github.com/UMCUGenetics/RNASeq-NF), with the appropriate reference genome for each on species. In short, reads were trimmed with TrimGalore v0.6.5 (Krueger et al. 2023) and rRNA was removed with SortMeRNA v4.3.3 (Kopylova, Noé, and Touzet 2012). Alignment was performed with STAR v2.7.3a (Dobin et al. 2013) and PCR duplicates were identified with Sambamba MarkDup v0.7.0 (Tarasov et al. 2015). Gene expression was quantificatied with featureCounts v2.0.0 (Liao, Smyth, and Shi 2014), generating RPKM values.

### Lettuce Expression Browser

The LEB hosts four transcriptome datasets, which are accessible via individual tabs. To support this modular structure, the LEB was built as an Rshiny app (Chang et al. 2025) within a Docker container, and is hosted online at https://lettuce.bioinformatics.nl. Each module was constructed in two steps: (1) preprocessing the transcriptomic data for fast access, and (2) visualising gene expression. Preprocessing involved converting RPKM values to TPM for each dataset and storing the resulting data as R objects (.RDS files) to allow rapid access. Metadata was parsed and the samples were grouped by experiment and species. To ensure data quality, potential mix-ups and low-quality samples (fewer than 1 million reads) were excluded, with mix-ups identified based on metadata inconsistencies and PCA anomalies. The final dataset includes the number of replicates, average expression levels and standard deviations, all stored in .RDS files for fast access.

Visualisation was implemented using the ggPlantmap R package (Jo and Kajala 2024), which overlays expression data onto schematic plant images using a colour gradient. Images were manually traced using ICY software (https://icy.bioimageanalysis.org) based on lettuce plant images from each experiment. The maps are stored in XML format for visualisation. By selecting a species/cultivar name and gene identifier, a plot is generated to display the organ-specific gene expression in TPM. The plot is made interactive with plotly, allowing for user interaction, and can be downloaded (Wickham 2016).

Additional gene information was added to the Lettuce Expression Browser to help interpretation of results and to translate findings to other relevant plant species (the same as used in (van Workum, Mehrem, et al. 2024)). First, we identified functional domains for the longest isoform of each gene using InterProScan v5.53-87.0 (Jones et al. 2014). Homology across plant species was determined using PanTools v4.3.2 using the longest isoform of a representative set of high-quality eudicot genomes (Jonkheer et al. 2022). All information is stored in tables, which are loaded into a Flask-based custom API that can be queried through a “Gene Information” page in the LEB. In addition to this functional and homology information, we provide a direct hyperlink to a JBrowse2 instance of the genome assembly (Diesh et al. 2023). Also, as some users may use transcript identifiers instead of gene identifiers, transcript identifiers are redirected to their respective gene identifiers. A page for *A. thaliana* ARAPORT11 identifiers exists as well (next to the three *Lactuca* spp. pages) to enable translation of *A. thaliana* to *Lactuca* via homology or functional domains. Using the same approach, links are created between all of the three *Lactuca* spp. There are hyperlinks from the LEB to the relevant “Gene Information” page and *vice versa*. This allows users to explore detailed information about the visualised gene, search for genes of interest and retrieve specific gene-expression data across organs and selected tissues, developmental stages and conditions.

The LEB was developed in close collaboration with the authors of the experimental data. However, since the LEB is meant as a resource for the global lettuce community, testing with potential users was performed based on standards from the field of visual analytics (see **Methods S1**).

## Data statement

Sequencing data for this study have been deposited in the Sequencing Read Archive (SRA) at NCBI under accession numbers PRJNA1347382 (https://www.ncbi.nlm.nih.gov/sra/PRJNA1347382; for the “Seedling abiotic stress”), PRJNA1347432 (https://www.ncbi.nlm.nih.gov/sra/PRJNA1347432; for the “Seedling far-red time series”), PRJNA1347570 (https://www.ncbi.nlm.nih.gov/sra/PRJNA1347570; for the “Shoot developmental stages”) and PRJNA1347639 (https://www.ncbi.nlm.nih.gov/sra/PRJNA1347639; for the “Seedling tissue map”). XML files underlying the ggPlantmap visualisations in the LEB have been deposited at data.4TU (https://doi.org/10.4121/8a593c82-89d4-41ee-8747-4bc7fb07d844).

## Funding

This publication is part of the LettuceKnow project (Perspective Program P19-17) which is (partly) financed by the Dutch Research Council (NWO; AES) and the breeding companies BASF, Bejo Zaden B.V., Limagrain, Enza Zaden Research & Development B.V., Rijk Zwaan Breeding B.V., Syngenta Seeds B.V., and Takii and Company Ltd.

## Acknowledgements

We would like to thank Christa Testerink for help with the experimental design and Ferdinand Kirsten, Xiaoyan Liu and Samara Almeida Landman for their help with sampling.

## Supporting Information

**Figure S1: Growth of different *Lactuca* genotypes in the “Seedling tissue map” experiment**

**Figure S2: Light spectrum profiles used in the “Seedling abiotic stress” experiment**

**Figure S3: Principal Component Analysis (PCA) of *L. saligna* and *L. virosa* samples across experimental datasets**

**Figure S4: Principal Component Analysis (PCA) of the “Seedling far-red time series” dataset**

**Figure S5: Fine-scale dissection and multi-species analysis reveal tissue-specific expression**

**Figure S6: Stomatal density increases across later-developing leaf positions in lettuce.**

**Table S1: Timing of shoot developmental stages (BBCH scale) in different genotypes**

**Table S2: RNA-seq metadata**

**Table S3: Dissection and pooling of tissues for the “Seedling tissue map”**

**Methods S1: User-testing of the Lettuce Expression Browser**

## Contributions

D.J.M.v.W., A.H., R.H., and S.S. contributed to conceptualization. D.J.M.v.W. led software development, data analysis, and writing of the original draft. A.H. contributed to data collection, methodology, and writing of the original draft. R.H. edited the manuscript. M.C.A., S.S.S., J.v.L., E.S.v.d.B., T.F., G.F.A., A.P., and D.L. contributed to data collection, with M.C.A. performing RNA isolation, F.F.M.M. running RNA-seq processing pipelines and D.L. managing data curation. R.H, S.S, M.G.M.A., M.P., R.O., R.P., G.v.d.A., and M.E.S. provided supervision. A.H., S.S.S., J.v.L., E.S.v.d.B., T.F., G.F.A., A.P., R.H., M.G.M.A., M.P., R.O., R.P., and M.E.S. contributed to methodology. All authors reviewed and approved the final manuscript.

## Notes

https://www.ncbi.nlm.nih.gov/sra/PRJNA1347382

https://www.ncbi.nlm.nih.gov/sra/PRJNA1347432

https://www.ncbi.nlm.nih.gov/sra/PRJNA1347570

https://www.ncbi.nlm.nih.gov/sra/PRJNA1347639

### Competing Interest Statement

The authors have declared no competing interest.

