## Supplementary Figures for "The Lettuce Expression Browser: from lab to LEB"

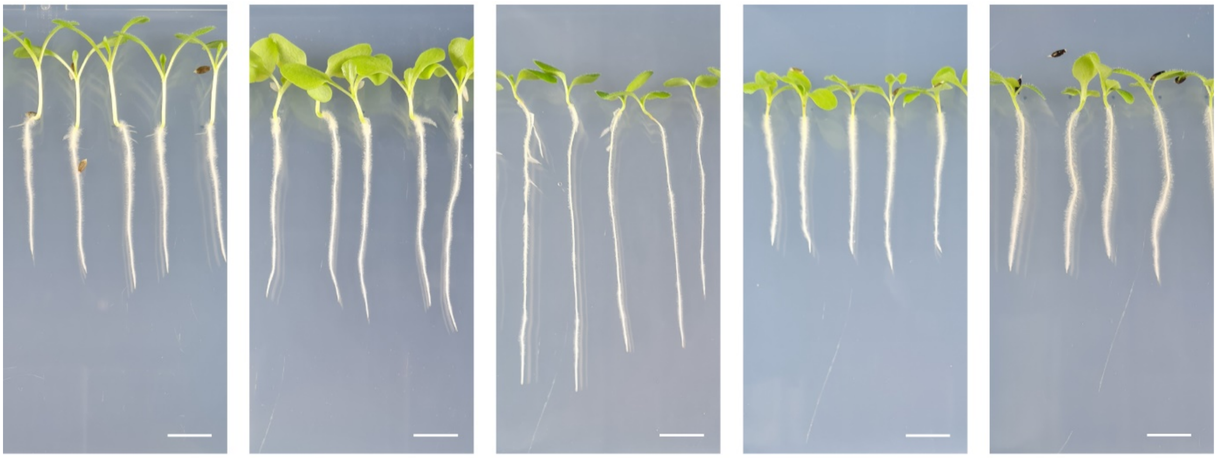

*L. sativa* 'Salinas' 7-day-old | *L. sativa* 'Cobham Green' 7-day-old | *L. serriola* 7-day-old | *L. saligna* 10-day-old | *L. virosa* 11-day-old

**Figure S1: Growth of different *Lactuca* genotypes in the "Seedling tissue map" experiment**
Seedlings of the five *Lactuca* genotypes were grown on ½MS plates for the indicated number of days to account for differences in growth rates. Scale bars, 1 cm.

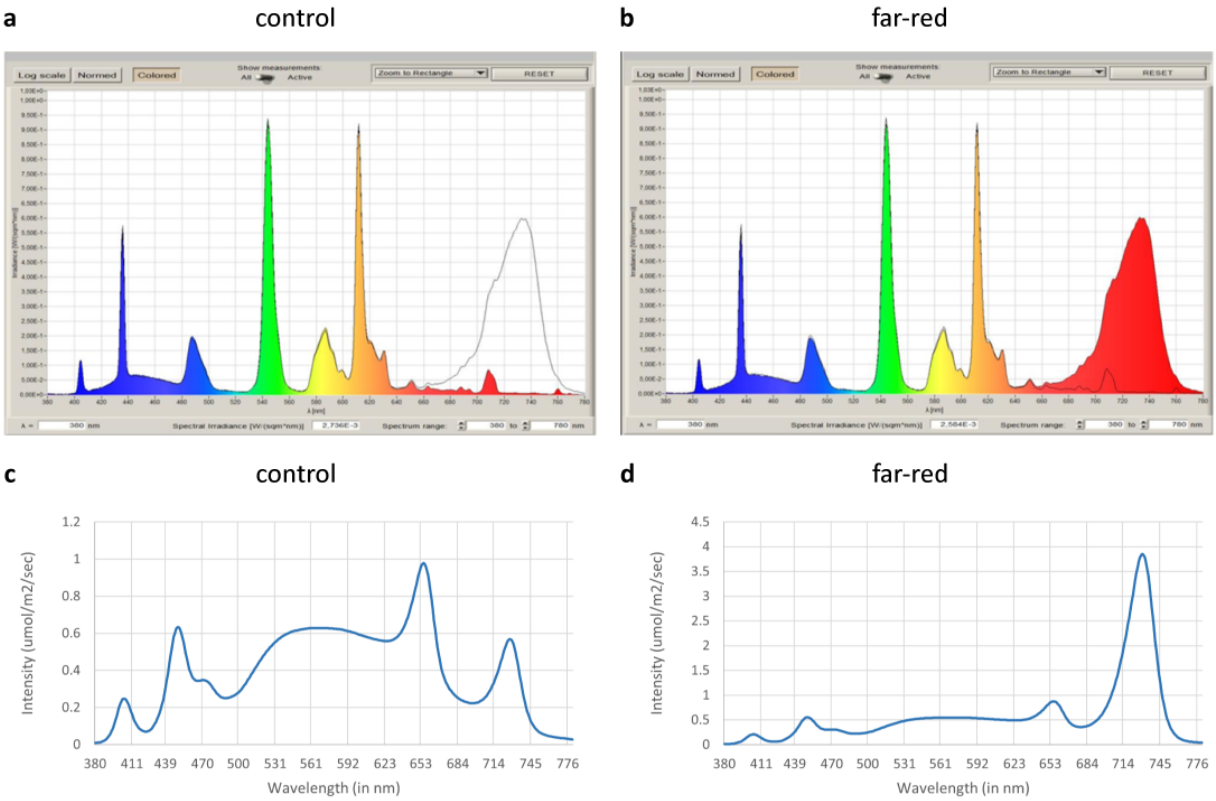


**Figure S2: Light spectrum profiles used in the experiments including far-red treatments**
**(a-b)** Light spectrum profiles used in the "Seedling abiotic stress" experiment in **(a)** control and salt stress treatments and **(b)** supplemental far-red light treatment. **(c-d)** Light spectrum profiles used in the "Seedling far-red time series" experiment in **(c)** control treatment and **(d)** supplemental far-red light treatment.

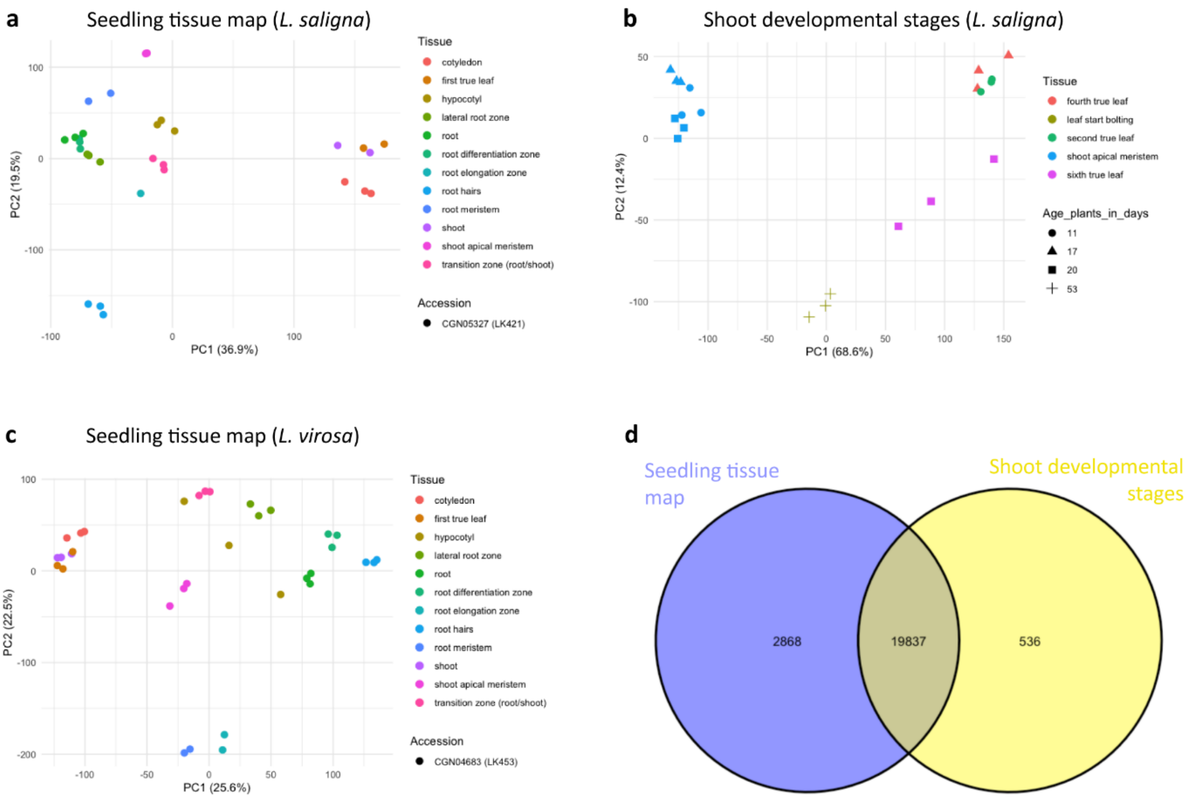


**Figure S3: Principal Component Analysis (PCA) of *L. saligna* and *L. virosa* samples across experimental datasets**

**(a-b)** PCA plots of the **(a)** “Seedling tissue map” and **(b)** “Shoot developmental stages” datasets, showing *L. saligna* samples, coloured by tissue. **(c)** PCA plot of the “Seedling tissue map” dataset, showing *L. virosa* samples, coloured by tissue. **(d)** Venn diagram depicting the number of genes expressed (TPM ≥ 2 in at least two samples) in each *L. saligna* dataset and their overlap across experiments.

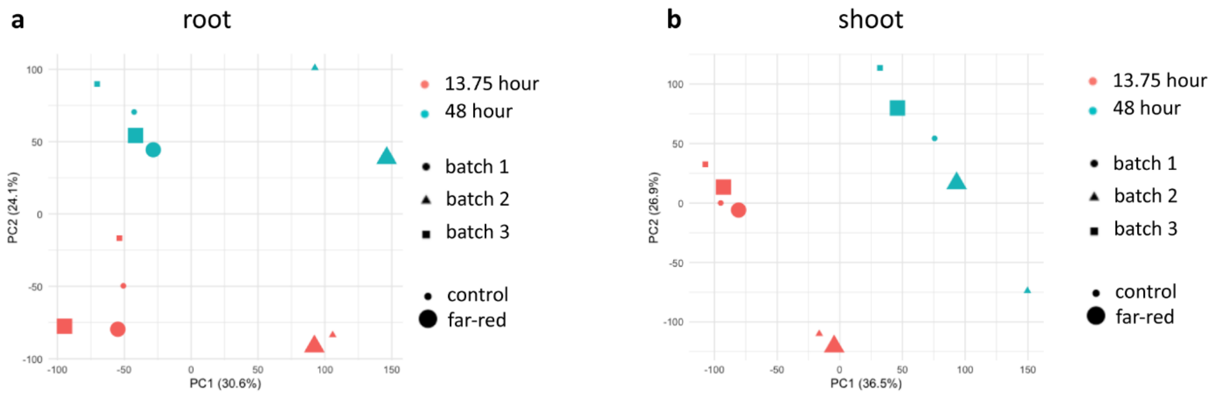


**Figure S4: Principal Component Analysis (PCA) of the "Seedling far-red time series" dataset (a-b)** PCA plots of the "Seedling far-red time series" dataset for **(a)** root and **(b)** shoot samples. Colours indicate time-of-day, shapes indicate the growth batch (as replicates were grown independently) and point size represents the treatment.

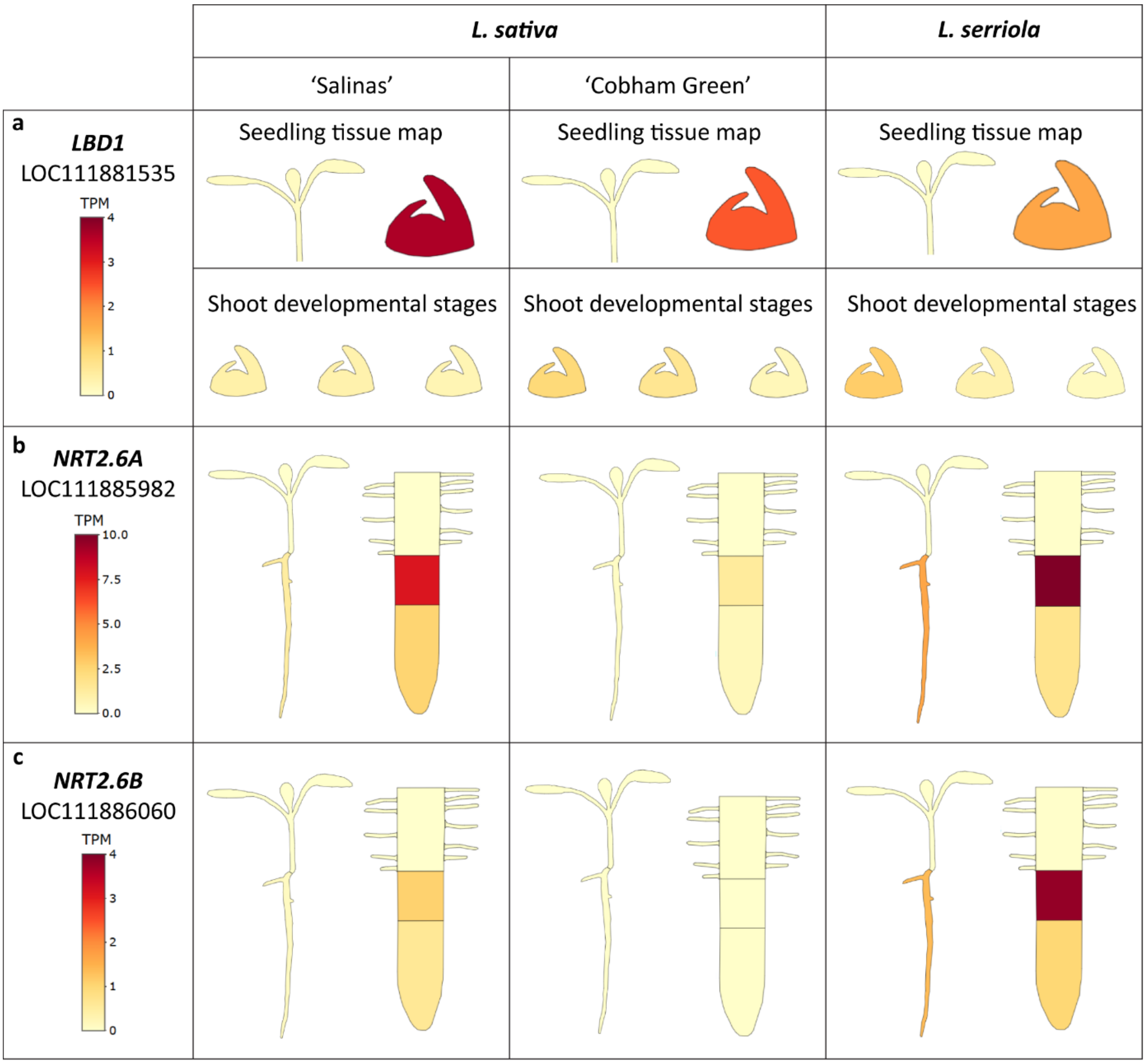


**Figure S5: Fine-scale dissection and multi-species analysis reveal tissue-specific expression**

**(a-c)** Expression of three genes that were expressed (TPM ≥ 2 in at least two samples) exclusively in the "Seedling tissue map" dataset. **(a)** Expression of *LOB DOMAIN-CONTAINING PROTEIN 1* (*LBD1*, LOC111881535) was detected in the dissected shoot apical meristems of *L. sativa* and *L. serriola* seedlings in the "Seedling tissue map" but not in the "Shoot developmental stages" dataset. **(b)** A homolog *of HIGH AFFINITY NITRATE TRANSPORTER 2.6* (*NRT2.6A*, LOC111885982) was specifically expressed in the root elongation zone of *L. sativa* 'Salinas' and in *L. serriola* seedlings. **(c)** Another *NRT2.6* homolog (*NRT2.6B*, LOC111886060) was found to be specifically expressed in the root elongation zone of *L. serriola* seedlings. TPM: transcripts per million.

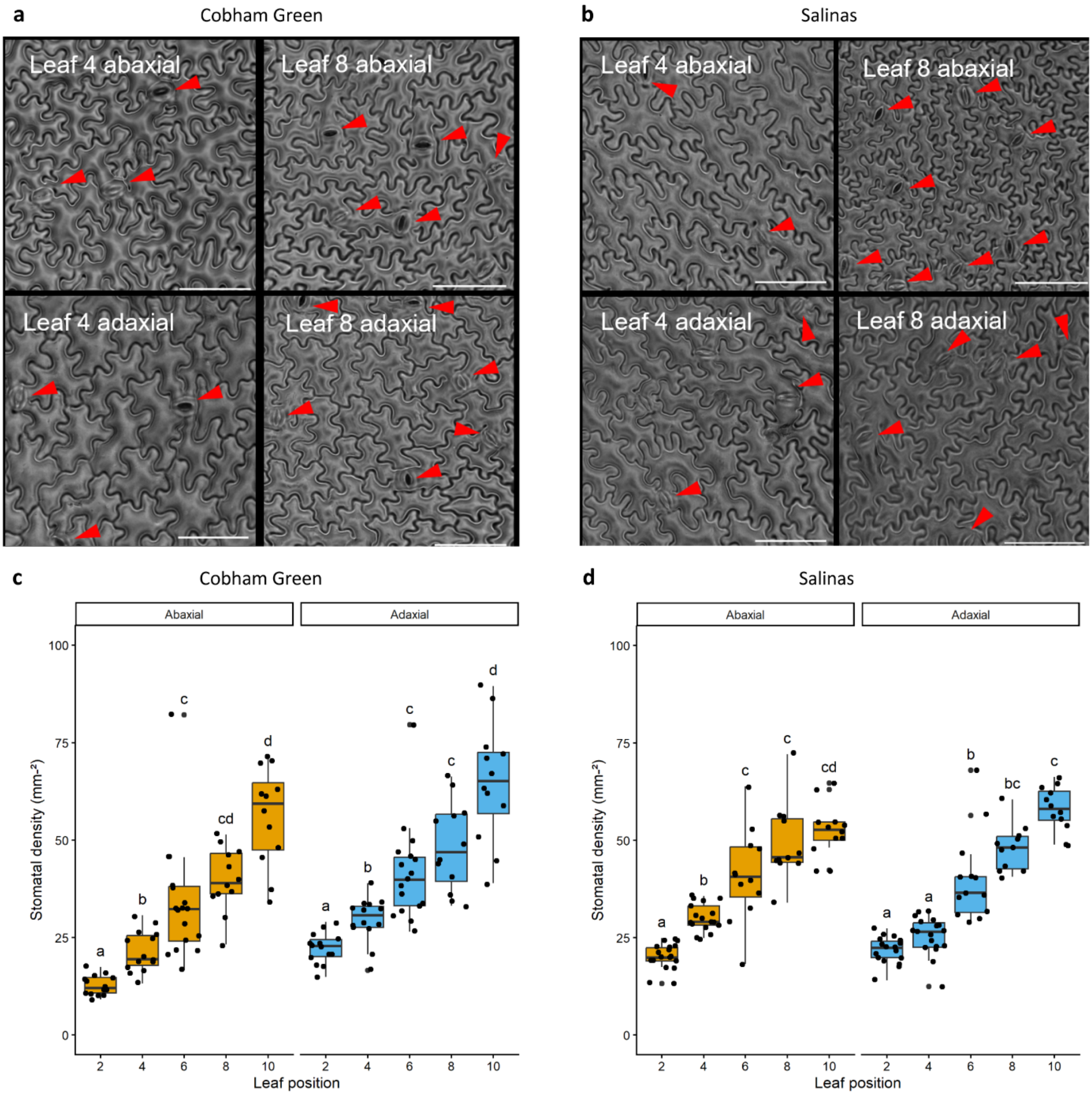


**Figure S6: Stomatal density increases across later-developing leaf positions in lettuce.** **(a-b)** Imprints of the abaxial and adaxial sides of fully expanded 4th and 8th leaves of 35-day-old *L. sativa* 'Cobham Green' **(a)** and 'Salinas' **(b)** plants. Stomata are indicated with red arrowheads. Scale bar = 100 µm. **(c-d)** Quantification of mean stomatal density in 'Cobham Green' **(c)** and 'Salinas' **(d)** leaves using images as shown in **(a-b)**. Significant differences, as determined using a one-way ANOVA followed by a post hoc Tukey's test, are indicated by different letters above boxplots ($p < 0.05$, $n > 10$).
