## Supplementary Methods for "The Lettuce Expression Browser: from lab to LEB"

**Methods S1: User-testing of the Lettuce Expression Browser**

**Introduction**

The Lettuce Expression Browser (LEB) was developed to support the global lettuce community. To ensure the tool is intuitive, usable and user-friendly, we spent considerable effort in user-testing throughout the development. The LEB was built through close collaboration between a bioinformatician and an experimental biologist of one of the four experiments in the LEB. After the initial design was complete and all four experiments had been integrated, we started a cycle of continuous user-testing and iterative refinement based on user feedback.

**Approach**

To standardise the user testing process, we first defined the core functionalities of the LEB, which formed the basis of task-based user evaluations. Participants were asked to complete specific tasks while thinking out loud, allowing us to observe the thought process. The four core functionalities were:

1. Retrieve expression values (TPM) for gene X under condition Y.
2. Determine whether gene X exhibits tissue- or treatment-specific expression.
3. Compare the expression profiles of gene X and gene Y.
4. Download expression plots and adjust their parameters if necessary.

For each functionality, we chose an example gene. For example, for functionality 2 we had two example genes: LOC111903971 (*STOMAGEN*, which is expressed in leaves) and LOC111891693 (*UBIQUITIN*, which is expressed everywhere).

User tests were conducted in stages, starting with those most familiar with the LEB. First, we performed a user test with the original authors of the LEB to align on task expectations. This was followed by user tests with the authors of the additional experiments in the LEB. A total of five initial user tests were performed, after which we summarised the main feedback and addressed recurring issues. We then expanded testing to include bioinformaticians (both academic and industrial) who had no prior involvement in LEB's development. Again, we summarised their main feedback and addressed recurring issues.

**Results**

*General impression*

In general, users very much appreciated the visualisation and interaction with the LEB. Some highlights from users were the interconnectedness of the different pages (from LEB to Gene Information and back), the clarity of the visualisations and the ease of use. The users all successfully performed the given assignments, indicating general usability for both experimental biologists and bioinformaticians. Although some bugs were reported during the user tests, they never stood in the way of using the LEB and all users found workarounds for the bugs they encountered. All users had very similar impressions of the LEB, with only one point of divergence: the amount of information. While some users appreciated the large amount of metadata and information about the genes they searched for, others found certain pages overwhelming.

*Reported bugs*

We tracked all reported bugs, issues and questions during the user testing phases, prioritising those that affected core functionality. During the five initial user tests (with those familiar to the LEB), one recurring issue was a malfunctioning gene selection drop-down menu. Since selecting a gene is a core functionality of the LEB (see functionality 1.), this was the first thing we improved in the LEB during the user tests. Another common problem was that there was no way to return to the tile page (the 'home' of lettuce.bioinformatics.nl) when the LEB was in full screen mode. To resolve this, we added a 'Back to main page' button in the LEB.

**Conclusion**

The user tests provided valuable insights in the interaction of the users with the LEB. As general understanding and interaction of the LEB by the users was as intended, we are confident about the usefulness of the LEB for the global lettuce community. The user tests also helped us assess what core functionalities were used most by users. Nevertheless, smaller bugs or issues related to non-core functionalities may still occur within the LEB, but these will be part of general maintenance of the web portal.
